# Evolutionary bioenergetics of sporulation

**DOI:** 10.1101/2025.08.26.672491

**Authors:** Canan Karakoç, William R. Shoemaker, Jay T. Lennon

**Affiliations:** Department of Biology, Indiana University, Bloomington, IN 47405, USA; School of Biological Sciences, Georgia Institute of Technology, Atlanta, GA 30332, USA; Quantitative Life Sciences, The Abdus Salam International Centre for Theoretical Physics (ICTP), Trieste 34151, Italy

**Keywords:** development, metabolism, ecology, dormancy, life history

## Abstract

Energy is required for the expression and maintenance of complex traits. In many habitats, however, free energy available to support biosynthesis is in vanishingly short supply. As a result, many taxa have evolved persistence strategies that support survival in unfavorable environments. Among these is sporulation, an ancient bacterial program governed by a large genetic network that requires energy for both regulation and execution. Yet sporulation is a last resort, initiated when cellular energy is nearly exhausted. To resolve this paradox, we quantified the energetic cost of sporulation in units of ATP by integrating time-resolved genome, transcriptome, and proteome profiles. The full cost of the spore cycle, including both formation and revival, ranks among the most energy-intensive processes in the bacterial cell, requiring almost 10^10^ ATP and consuming about 10% of the total energy budget. The majority of this cost arises from translation, membrane synthesis, and protein turnover. Despite its considerable upfront investment, sporulation enables long-term survival and becomes optimal when harsh conditions extend over timescales of months or longer. This trade-off between immediate cost and delayed benefit helps explain when sporulation is maintained or replaced by alternative strategies. By incorporating our estimates into mechanistic models, we show how metabolic constraints shape sporulation efficiency, while genome-wide mutation accumulation data reveal that even modest energetic burdens can become visible to selection, influencing the evolutionary fate of this complex and widespread trait.

---

Life requires energy to support growth, homeostasis, and long-term persistence. Energy is stored and transferred by molecules such as adenosine triphosphate (ATP), which fuel catabolic and anabolic reactions that govern basal metabolism, biosynthesis, and cellular repair [29, 61]. These energy carriers also power gene regulation and translation, processes essential for building and maintaining cellular structures and organismal traits. In the early history of life, energetic constraints shaped the emergence of self-replicating systems and the evolution of core molecular functions [34]. Today, the availability of energy continues to influence both physiological and developmental processes, from relatively simple biochemical pathways to more elaborate features such as secretion systems, sensory networks, and multicellular differentiation [41, 58, 78]. While it is generally assumed that traits are maintained when fitness benefits outweigh their energetic costs, such trade-offs are rarely quantified, particularly under conditions that limit metabolism [73, 80]. As such, the role of bioenergetics in determining the evolutionary fate of complex traits remains largely unresolved.

In nature, organisms often inhabit environments where energy is insufficient to support sustained growth and reproduction [36]. To endure these unfavorable conditions, many species have evolved mechanisms of persistence that buffer against demographic variance and environmental stochasticity [84]. Dormancy is one such strategy, whereby individuals to enter a reversible state of reduced metabolic activity. In some cases, dormancy arises passively as metabolism slows in response to environmental stress. In others, it is complex and actively regulated, requiring precise gene expression and cellular remodeling [35]. One of the most elaborate and well-characterized forms of dormancy is endospore formation, which is found among bacteria like *Bacillus* and *Clostridia*. Endospores are among the most metabolically inert and long-lived biological entities [70]. They are remarkably resistant to harsh environmental conditions, including elevated temperatures, high doses of ionizing radiation, extreme energy limitation, and the vacuum of space [55, 69]. Consequently, endospores are widely distributed across environmental, engineered, and host-associated ecosystems. With an estimated 10^28^ individuals in marine sediments alone [88], endospores rank among the most abundant cell types on Earth.

Despite its protective benefits, sporulation comes with an energetic cost. Building a spore demands sustained investment of cellular resources over a prolonged developmental timeline. Roughly, 5% of the *Bacillus subtilis* genome is devoted to this process, encoding genes involved in the synthesis of spore structures (e.g., cortex and coat proteins), DNA packaging and repair, and signaling pathways that guide cell fate decisions [22]. Sporulation is orchestrated by a precisely timed cascade of sigma factors, a family of proteins that enable RNA polymerase to bind specific gene promoters and initiate transcription [75, 83]. Each sigma factor is activated in one of two cellular compartments, directing stage-specific gene expression during development [83]. This tightly regulated sequence is punctuated by checkpoints that ensure the mother cell adequately provisions the developing forespore [83]. However, sporulation alone does not guarantee evolutionary success. To realize a fitness benefit, the resting cell must ultimately be revived through the processes of germination and outgrowth. This developmental transition is regulated in part by changes in electrochemical potential [30]. The resulting ion gradients serve as a memory-like mechanism for evaluating environmental quality and decision making. Although germination and outgrowth are faster than sporulation, they entail a complete turnover of the proteome [77], adding further costs to the full dormancy cycle.

The energetic demands of spore formation and revival have important implications for the evolution of microbial populations. An ancestral trait within the phylum Bacillota, sporulation is thought to have originated nearly three billion years ago [4]. Over this geological timespan, it has been conserved in some groups, partially retained in others, and entirely eliminated in major clades such as *Staphylococcaceae* and *Lactobacillaceae*, with consequences for molecular evolutionary dynamics [15]. For example, phylogenomic comparisons reveal that non-spore-forming bacteria diversify more rapidly than their spore-forming counterparts, while experimental evolution shows that the loss of sporulation ultimately reduces genetic diversity within populations [74]. The evolutionary loss of sporulation may occur predominantly through neutral processes, as the underlying genetic network represents a large mutational target. Models predict that, in the absence of selection, this trait could decay within approximately 10^8^ generations [47]. Although some argue that the genetic burden of maintaining sporulation machinery should be minimal, the cost of unused genes can still be visible to selection. Indeed, spore loss has been associated with genome streamlining, in which deletions in non-expressed genes are favored [14], potentially reflecting energetic cost savings. However, empirical evidence supporting either neutral or selective explanations remains limited and inconclusive [49].

Accurately quantifying the energetic costs of complex traits may clarify the conditions that favor their maintenance or loss. However, standard measurements of energy demand, such as oxygen consumption, heat production, or metabolite turnover, often lack precision, use units that are difficult to interpret in cellular terms, and are challenging to compare across biological scales. To address these limitations, we used a quantitative bioenergetic framework grounded in ATP equivalents, the universal currency of energy transfer in all forms of life [43]. By integrating genomic information with temporally resolved transcriptomic and proteomic measurements, we estimated the energetic demands of sporulation and germination. This approach enabled us to quantify both opportunity costs (*P*_*O*_) and direct costs (*P*_*D*_) associated with precursor synthesis, gene expression, and proteome turnover across developmental time. With these estimates, we compared the energetic costs of dormancy to other components of the total cellular budget, including baseline energy demands and other persistence traits that may confer advantages under fluctuating or suboptimal conditions. Finally, we incorporated the estimated costs into a mechanistic model to examine how energetic constraints influence sporulation efficiency. By combining these results with genome-scale mutation accumulation data, we tested predictions about how bioenergetics shape the relative contribution of neutral and selective processes to the long-term maintenance of sporulation.

## Results and Discussion

We begin with a detailed accounting of the energetic costs underlying the spore life cycle of the model bacterium *Bacillus subtilis*, resolving patterns of energy expenditure over time, across developmental transitions, and within subcellular compartments. Our analysis shows that the complete spore life cycle demands a major energetic investment, consuming about 10% of the total cellular energy budget. Most of this cost arises from opportunity costs (*P*_*O*_), including the diversion of metabolic precursors, along with the direct synthesis of spore-specific macromolecules. By incorporating these estimates into a mechanistic model, we identify thresholds beyond which dormancy is no longer advantageous and demonstrate how energetic constraints limit sporulation efficiency, defined as the proportion of cells that complete the transition to an endospore. Based on genome-scale comparisons, our findings suggest that positive selection driven by energetic costs contributes to the evolutionary loss of sporulation, alongside mutational decay through neutral processes. The bioenergetic framework developed here provides a conceptual and quantitative foundation for understanding how resource allocation strategies shape the ecological and evolutionary dynamics of complex traits.

### 1. Energetics of spore formation

We estimate the total cost of producing a single spore to be *∼* 2.4 *×* 10^9^ ATP (*P*_*T*_). The majority of this cost (80%) is attributable to opportunity costs (*P*_*O*_), which reflect the diversion of precursor metabolites away from growth. The remaining 20% consists of direct costs (*P*_*D*_), representing ATP hydrolysis events required for the biosynthesis and polymerization of molecular building blocks. Across both *P*_*O*_ and *P*_*D*_, we found that translation accounts for the largest share of energy expenditure (68%), followed by genome replication (17%), and transcription (3%). Upon initiation of sporulation, the cell must duplicate its entire genome [79]. This requires 4 × 10^8^ *P*_*T*_, on the same order as the 7.8 × 10^7^ *P*_*T*_ needed to transcribe spore-related genes. Sporulation genes alone make up ∼ 21% of the *Bacillus subtilis* genome and contribute 18% (∼7.2 × 10^7^ *P*_*T*_) of the total replication cost. Although membrane synthesis has been hypothesized to impose a substantial energetic burden on cells [44], we estimate that the lipid required for the 1 *µm* septum that separates the mother cell from the forespore only amounts to 12% of *P*_*T*_ incurred during sporulation.

Our findings reveal that spore formation is characterized by a pronounced asymmetry in energetic investment, with the mother cell bearing the majority of the biosynthetic burden. We estimate that the mother cell accounts for roughly 87% of the total costs, while the forespore contributes the remaining 13%. Approximately 67% of the total energy expenditure, including the cost of genome replication, occurs within the first hour of development. This early demand drops off approximately exponentially as development progresses, a pattern reminiscent of maternal investment during animal embryogenesis [17]. Following initiation, septum formation creates two unequally sized cellular compartments, establishing the physical basis for asymmetric investment. More than one hundred genes are activated within the forespore to support differentiation and maintain coordination with the mother cell [83]. Throughout development, the mother cell delivers precursors and recycled building blocks to the forespore through a tubular intercellular channel [7, 21], reducing the need for *de novo* biosynthesis. This structure also mediates continued signaling between compartments during intermediate and late stages of development [25]. Ultimately, the mother cell lyses, releasing the mature spore into the environment (Fig. 1).

**Figure 1.**
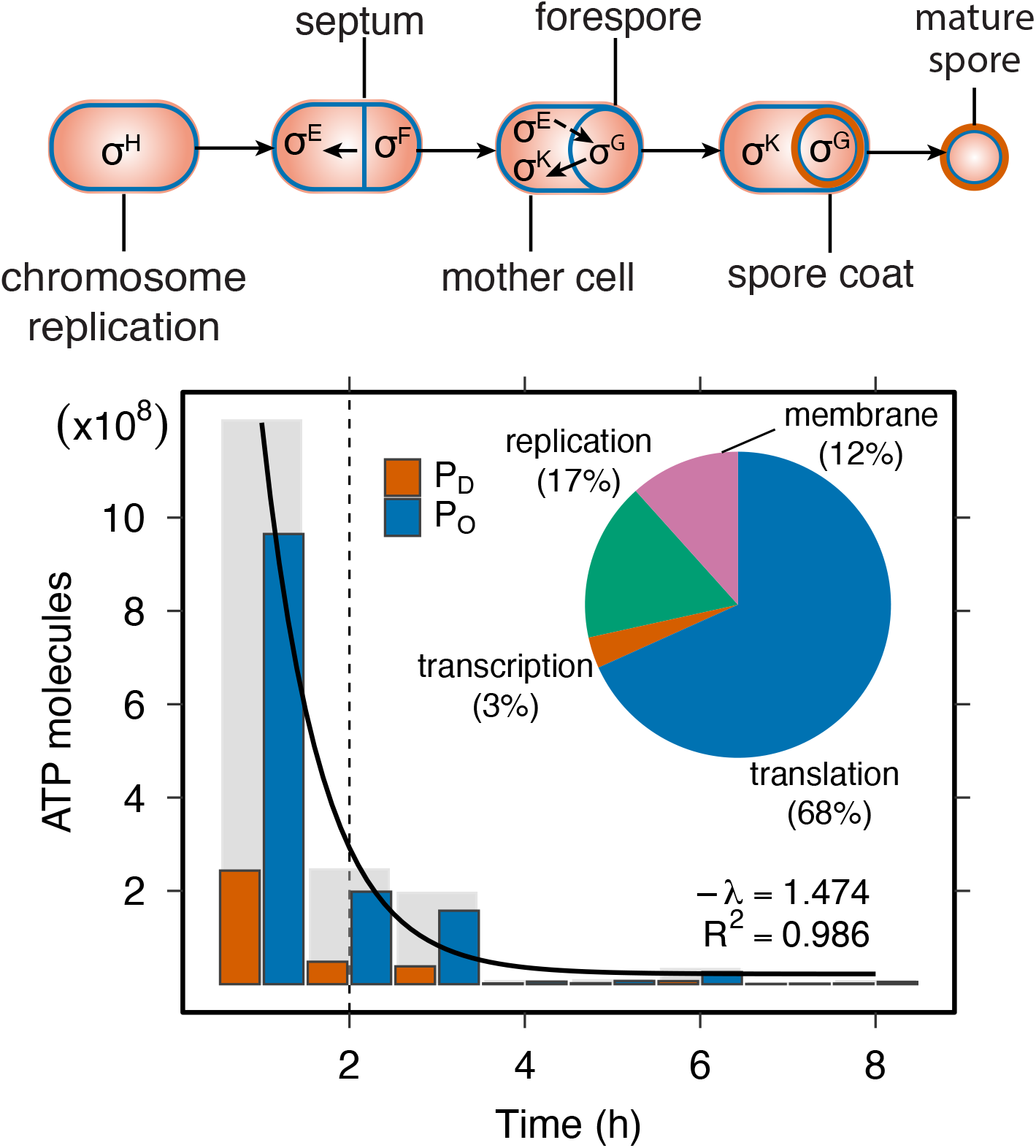
Energetic cost of spore formation. Top: Major stages of endospore development. Bottom: Energetic cost of spore formation over time, measured in ATP molecules, and includes transcriptional and translational investments. Total costs (*P*_*T*_, gray bars) comprise opportunity costs (*P*_*O*_, blue bars) linked to biosynthesis and direct costs (*P*_*D*_, orange bars) associated with polymerization. The costs of spore formation decline approximately exponentially over time. The embedded pie chart illustrates the proportional contribution (% of *P*_*T*_) associated with key processes, including replication of the whole genome and septum synthesis, along with transcription and translation.

Spore formation poses a risk for cells navigating unpredictable environments. Development unfolds slowly, requiring 8-10 h to complete compared to the 30-minute to 1-hour division cycle that is typical during vegetative growth. As a result, cells may engage in a costly process that offers no benefit if conditions improve. Our accounting shows that by the time the cell reaches the commitment point, ∼2 h after initiation, 85% of the total transcriptional and translational costs of sporulation have already been incurred. If precursor molecules are fully recycled, then only 20% of these costs are truly non-recoverable (i.e., we charge polymerization and activation in *P*_*D*_, but credit full recycling in *P*_*O*_). Nonetheless, this early phase still represents a substantial upfront investment. Prior to this checkpoint, cells can reverse course in response to environmental cues, although doing so results in a partial loss of energy [25]. Once SpoIIE activates asymmetric septum formation and *σ*_F_ is engaged [24], development becomes irreversible (Fig. 1). Beyond this threshold, the cell continues to invest heavily in spore-specific proteins, including structural components of the coat, small acid-soluble proteins that protect DNA from damage, and enzymes such as proteases that are required for revival [23, 66, 68].

### 2. Energetics of spore revival

Spore revival, which includes germination and subsequent outgrowth, is essential to the success of dormancy. Our analyses reveal that revival is even more energetically costly than spore formation, with total expenditures exceeding 6.8 10^9^ *P*_*T*_ (Fig. 2). Although germination (15 min) and outgrowth (3.5 h) proceed more rapidly than sporulation, they demand significant energy to reestablish vegetative growth.

**Figure 2.**
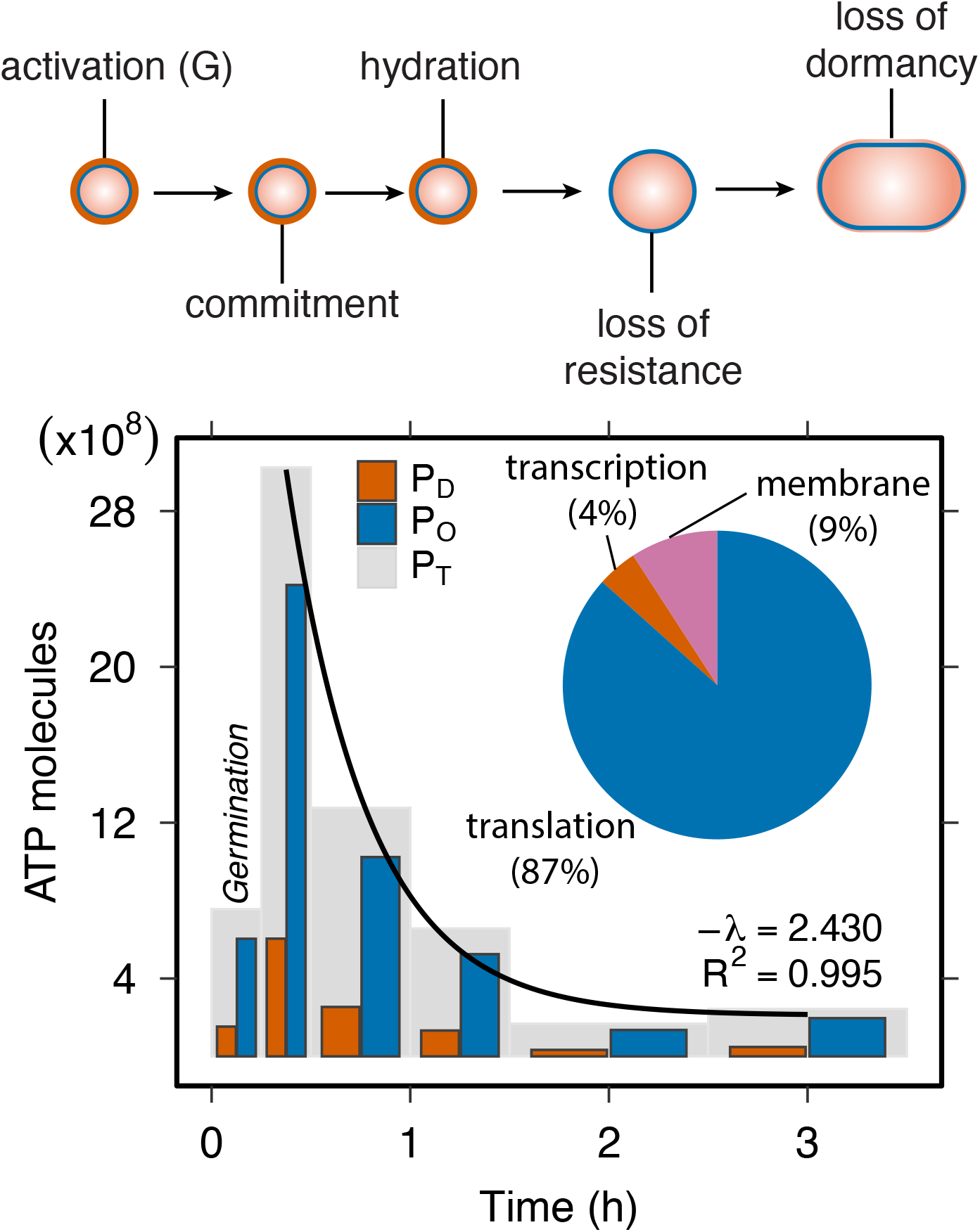
Energetic cost of spore revival. Top: Major stages of spore revival, which include germination followed by outgrowth. Bottom: Energetic costs of transcription and translation over time, measured in ATP molecules based on protein expression. Total costs (*P*_*T*_, gray bars) comprise opportunity costs (*P*_*O*_, blue bars) linked to biosynthesis and direct costs (*P*_*D*_, orange bars) associated with polymerization. Revival costs decline approximately exponentially over time. The embedded pie chart illustrates the proportional contribution (% of *P*_*T*_) of key processes, including transcription, translation, and membrane synthesis.

During the initial 15 min of revival, transcription and translation require ∼7.6 × 10^8^ *P*_*T*_ (1.5 × 10^8^ *P*_*D*_ and 6 × 10^8^ *P*_*O*_), representing 12% of the total revival cost. In contrast, the outgrowth phase consumes the vast majority (88%) of the energy: ∼5.4 × 10^9^ *P*_*T*_ for transcription and translation alone (1.1 × 10^9^ *P*_*D*_ and 4.3 × 10^9^ *P*_*O*_), and an additional ∼6.9 × 10^8^ *P*_*T*_ for membrane biogenesis (5.4 × 10^7^ *P*_*D*_ and 6.4 × 10^8^ *P*_*O*_), even after accounting for recycling of 1/6 of spore membrane lipids [3, 62], considering that only 3% of the total membrane is synthesized during this period. These results reveal a highly uneven allocation of energy across core molecular processes during revival, with transcription accounting for just 4%, translation for 86%, and membrane assembly for 10% of total energy expenditure (Fig. 2). Overall, the high rate of energy expenditure during revival reflects the rapid reactivation of biosynthetic pathways and the need to rebuild a complete proteome, in contrast to sporulation, which relies on a more selective set of proteins. Although revival-specific genes contribute an estimated ∼4.6 × 10^7^ (*P*_*T*_), equivalent to approximately 11% of the energy required for genome replication, this cost is paid in advance at the onset of sporulation.

Despite their ability to rapidly reawaken, dormant spores contain remarkably little internal energy. Our bioenergetic accounting shows that germination alone requires ∼7.6 × 10^8^ *P*_*T*_. ATP and GTP concentrations within *Bacillus* spores are extremely low, at around 2 nmol/g, as estimated using magnetic resonance spectroscopy with ^31^P NMR [16]. This corresponds to roughly 10^2^ molecules per spore. Additional reserves including AMP (∼400 nmol/g) and ADP (∼100 nmol/g) [71], are also present in low concentrations. Assuming a spore mass of ∼200 femtograms [8], total endogenous stores amount to just 2 × 10^3^ ATP equivalents. These estimates reveal that the energy required for germination exceeds internal reserves by nearly five orders of magnitude. While endogenous pools may be sufficient for minimal maintenance or for initiating the early steps of germination, they are far from adequate to support the full transition from dormancy to active growth.

The apparent energy shortfall presents a fundamental problem concerning how a dormant spore generates enough ATP to complete germination. Our analysis suggests that spores overcome this limitation by mobilizing pre-packaged molecular reserves that are metabolized immediately upon rehydration. Within 5 min of germination onset, essential components such as enzymes, ribosomes, amino acids, and nucleotides are activated to initiate core metabolic processes [71]. A major source of amino acids is the rapid degradation of small acid-soluble proteins, which account for 10–20% of the spore’s proteome [67]. In addition, stored carbon sources, including 3-phosphoglyceric acid (∼2,700 nmol/g) and malate (∼3,000 nmol/g), fuel glycolysis and related metabolic pathways, yielding ∼2.1 × 10^8^ ATP in total [16, 50]. These reserves are substantial but insufficient to cover the full germination demand; they likely bridge the earliest reactivation steps until exogenous carbon is taken up during outgrowth. However, full outgrowth and the return to vegetative growth require additional energy from the external environment. Increased glucose uptake during outgrowth likely supports the elevated ATP production needed for biosynthesis and cell expansion [28]. Because both spore formation and revival impose substantial energetic costs, their evolutionary persistence must be evaluated relative to alternative cellular stress responses, which we explore in the next section.

### 3. Head-to-head bioenergetics of microbial survival

Spore-forming bacteria like *Bacillus* use a tiered set of stress responses that differ in reversibility, energy cost, and long-term benefits. The earliest responses, coordinated by the alternative sigma factor *σ*_B_, involve low-cost physiological adjustments such as DNA repair, redox homeostasis, and osmoprotection [59, 64]. These transient changes help cells buffer short-term environmental fluctuations without altering their developmental trajectory. If conditions worsen, cells may adopt facultative strategies such as motility [9], cannibalism [26], oligotrophic survival [20], or competence [12], which offer moderate costs and reversible outcomes.

When other strategies fail, cells engage in spore formation, a developmental shift that requires a major upfront investment but offers long-term protection. We estimate that the complete spore cycle, including formation and revival, requires ∼9.2 × 10^9^ ATP, or about 10% of the total cellular energy budget (∼9.4 × 10^10^). This makes the spore life cycle one of the most energy-intensive programs in the cell, far exceeding the cost of short-term stress responses (Fig. 3). These findings align with recent estimates highlighting the energetic burden of structural and developmental processes [89].

**Figure 3.**
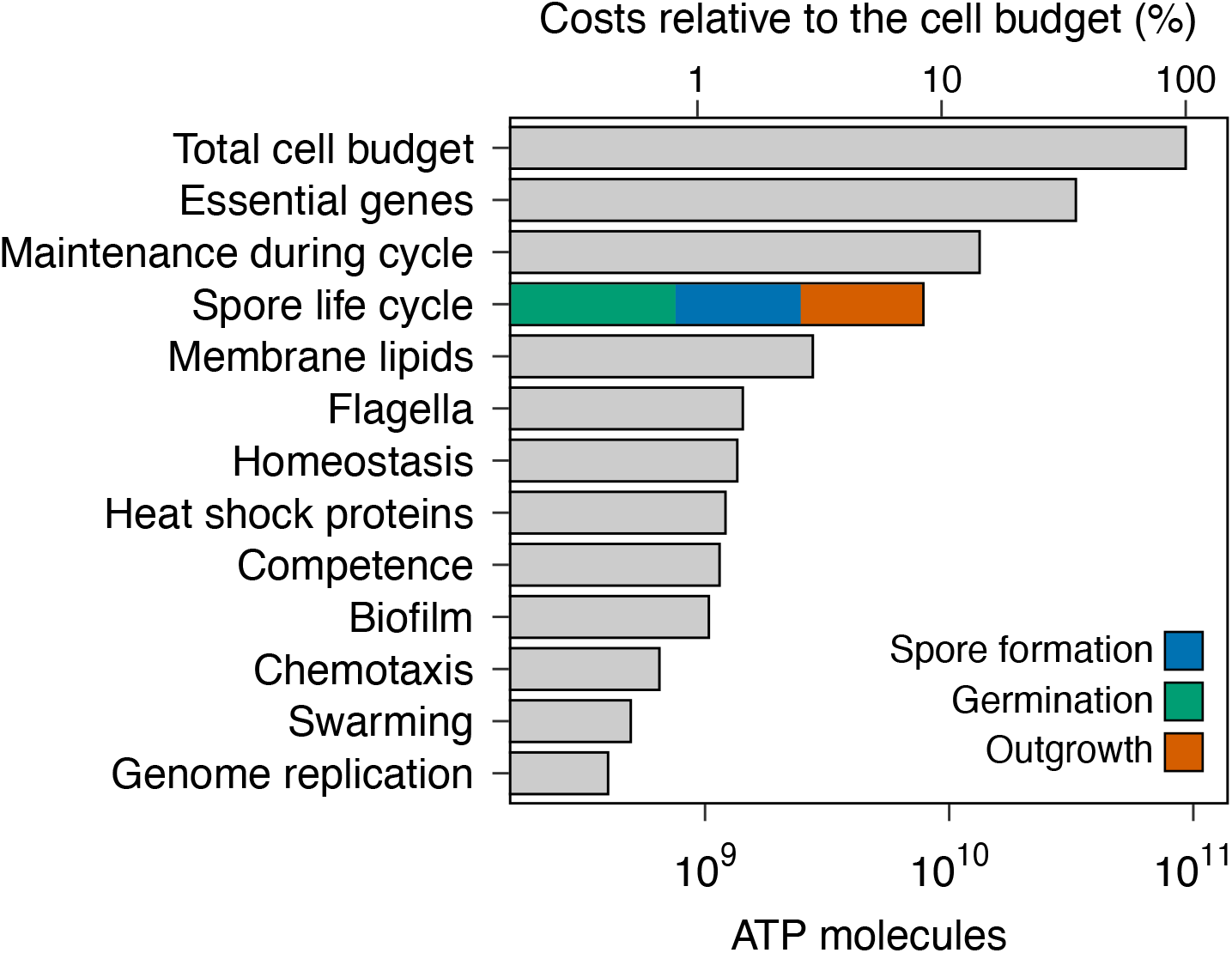
Comparative costs of spore life cycle and other cellular processes. Bars show build-only costs per cell (ATP molecules) for non-developmental traits (transcription and translation needed to build the mRNAs and proteins once), with no maintenance or degradation. Two global references are included: Genome replication (whole-genome DNA duplication) and membrane lipids (whole-cell bilayer, protein-occlusion corrected). The spore life cycle bar aggregates spore formation, germination, and outgrowth (transcription and translation only); indicated with colored cumulative phase totals. The maintenance during the cycle bar equals the per-hour maintenance cost multiplied by the program window (8 h spore formation, 0.25 h germination, 3.25 h outgrowth). The total cell budget is the per-generation growth/build cost plus one generation of maintenance (here, growth and maintenance for about 1 hour at 20°C, from [43]). The top (secondary) x-axis expresses positions as % of the total cell budget. All other trait bars exclude genome replication and membrane synthesis to avoid double-counting.

Spore formation is a front-loaded strategy whose benefits increase over time. Using a basal maintenance rate of *∼* 1.2 *×* 10^9^ ATP h^*−*1^ at 20°C, the break-even time is

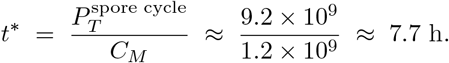

If cells down-shift maintenance by 10 (∼1.2 × 10^8^ ATP h^*−*1^), *t*^*∗*^ *≈* 77 h (∼3.2 d); with a 100× reduction (∼1.2 × 10^7^ ATP h^*−*1^), *t*^*∗*^ *≈* 770 h (∼32 d). Thus, without committing to sporulation, *Bacillus* can remain viable for extended periods at low power draw [20], but under prolonged or unpredictable stress the upfront cost of dormancy becomes the energetically optimal investment. Note that maintenance and growth conditions change at different temperatures, so these numbers will change accordingly.

Although commonly viewed as a last resort, sporulation can represent a bioenergetically optimal strategy under prolonged or unpredictable stress. Such conditions are widespread in nature, including energy-limited soils and sediments where resource inputs are both variable and vanishingly small [36]. By comparing the energy costs of microbial stress responses, our analysis reframes dormancy not as a passive fallback but as a strategic investment shaped by ecological, evolutionary, and developmental constraints.

## 4. Individual costs to collective outcomes: sporulation efficiency

Understanding the energetic costs of spore formation and revival has the potential to explain how dormancy strategies scale up to shape population-level phenomena. Excluding transitional states, an individual *Bacillus* exists either as metabolically active vegetative cell or as dormant spore (Figs. 1, 2). Yet this binary classification does not lead to uniform behavior across the population. Instead, the fraction of individuals that undergo sporulation, known as sporulation efficiency, is highly variable. In batch culture, where rapid physiological shifts and unbounded population growth eventually lead to resource exhaustion, the median sporulation efficiency is ∼30% but can range from 0 to 100% (Fig. 4A). This variability is not limited to transient conditions. Even in populations subjected to long-term energy limitation lasting over 1,000 days, sporulation remains incomplete, again hovering around 30% [72]. Such patterns are often attributed to cellular decision-making or bet-hedging, but are less frequently viewed through the lens of energetic constraints.

**Figure 4.**
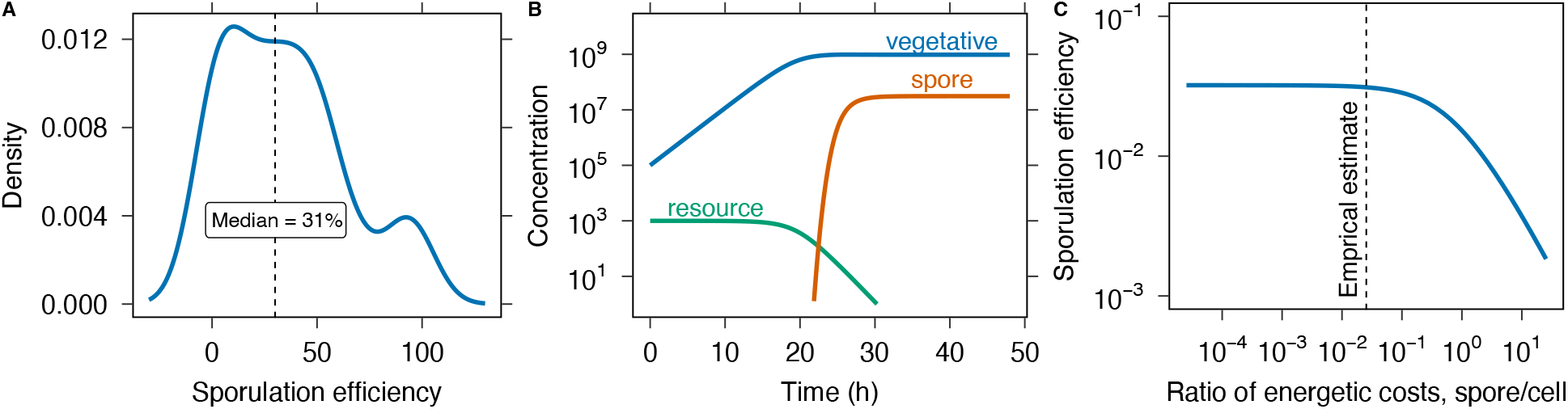
Energetic constraints on sporulation efficiency. A) The frequency of sporulation efficiency (fraction of cells that sporulate in a population) derived from publicly available data (Supporting Information, Table S2). B) In a dynamical model, resource depletion reduces cell growth and triggers spore formation. C) An increase in spore costs relative to cell costs results in a monotonic decline in sporulation efficiency. The dashed line represents the empirical estimate of the spore-to-cell cost ratio. The sharp decrease in efficiency as costs rise suggests that sporulation becomes energetically unfavorable with increasing costs.

Because all cellular features require energy to build and maintain, we examined how our estimates of energetic costs influence sporulation efficiency. To investigate this relationship, we developed a population dynamics model in an environment where resources decline over time:

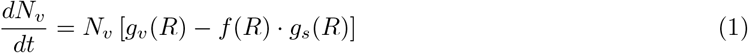

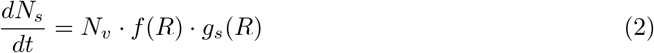

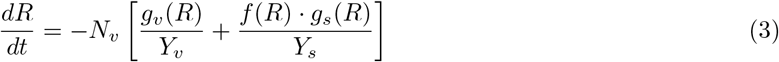

Here, *N*_*v*_ and *N*_*s*_ represent the concentrations of vegetative cells and spores, respectively. The functions *g*_*v*_(*R*), *g*_*s*_(*R*), and *f* (*R*) represent the per capita rates of cellular growth, sporulation, and the initiation of sporulation, each dependent on the concentration of available resources *R*. The yields *Y*_*v*_ and *Y*_*s*_ describe the efficiency of resource use for producing cells and spores, respectively, with ATP used as the unit of energetic currency (See Methods and Supporting Information for more detail).

Our model predicts that sporulation efficiency declines sharply when energetic costs exceed empirically derived estimates (Fig. 4C). This outcome reflects a trade-off between resource availability and the energy required for spore formation, which limits the number of cells that can successfully sporulate. By integrating empirical measurements with mechanistic modeling, we show that the relative cost of producing a spore compared to a vegetative cell can constrain the success of dormancy as a survival strategy.

## 5. Evolutionary maintenance of an energetically costly trait

Sporulation is an ancient survival strategy that originated billions of years ago. It confers resistance to harsh conditions, enables long-distance dispersal, and protects against viral infection [55, 65, 85]. These advantages have contributed to the prevalence of spore-forming lineages across the globe. Yet despite its benefits, this complex form of dormancy remains susceptible to evolutionary decay. Repeated and independent losses of sporulation have occurred across the Bacillota phylogeny [49, 74], often attributed to relaxed selection in environments that favor continuous growth [49]. In such settings, sporulation genes may no longer provide a fitness benefit and become prone to mutational degradation [48]. In addition to neutral decay, the energetic burden of unused genetic material may drive the adaptive deletion of sporulation loci, as even non-expressed genes and regulatory elements can impose fitness costs [27]. The relative importance of neutral versus selective forces in the loss of sporulation, however, remains unresolved.

When growth is favored, sporulation genes are typically not expressed, rendering their transcriptional and translational costs negligible. However, DNA replication still requires energy and resources. If even small costs are visible to selection, sporulation genes may be lost through the fixation of beneficial deletions. This contrasts with trait loss driven by neutral processes, in which mutations accumulate gradually through nucleotide substitutions. In *Bacillus*, the estimated rate of deletions is 1.2 × 10^*−*10^ per site per generation, while the rate of nucleotide substitution is 3.35 × 10^*−*10^ per site per generation, making substitutions nearly three times more frequent [76]. Whether the energetic cost of maintaining unused sporulation genes is sufficient to favor deletions over neutral decay depends on the strength of selection and the size of deletion. To examine this, we compared the fixation rate of a beneficial deletion to that of a neutral substitution:

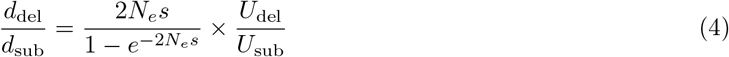

where *N*_*e*_ is the effective population size, *s* is the selection coefficient for a mutation, *U*_*del*_ is the rate of deletions, and *U*_*sub*_ is the rate of substitutions.

To examine how the relative fixation rate depends on deletion size, we combined our energetic cost estimates with published values for *N*_*e*_ and the empirical distribution of deletion sizes from mutation accumulation experiments [5, 76] (Fig.5A). Although deletions occur less frequently than substitutions, selection acting on large deletions (∼1kb) is strong enough to outweigh the rate of neutral substitutions. This length corresponds to the typical size of a gene, providing evidence that selection can drive the loss of sporulation in bacterial populations (Fig.5B). While previous studies have emphasized mutational degradation under relaxed selection [49], our analysis shows that the energetic burden of maintaining unused sporulation genes can be sufficient to favor their adaptive deletion. Even small metabolic costs, if persistent, may become visible to selection [47, 53]. Observations from host-associated Bacillota, where sporulation loss often coincides with genome streamlining and metabolic specialization, support the idea that energy-driven processes shape trait loss in natural populations under real-world conditions [6, 18].

**Figure 5.**
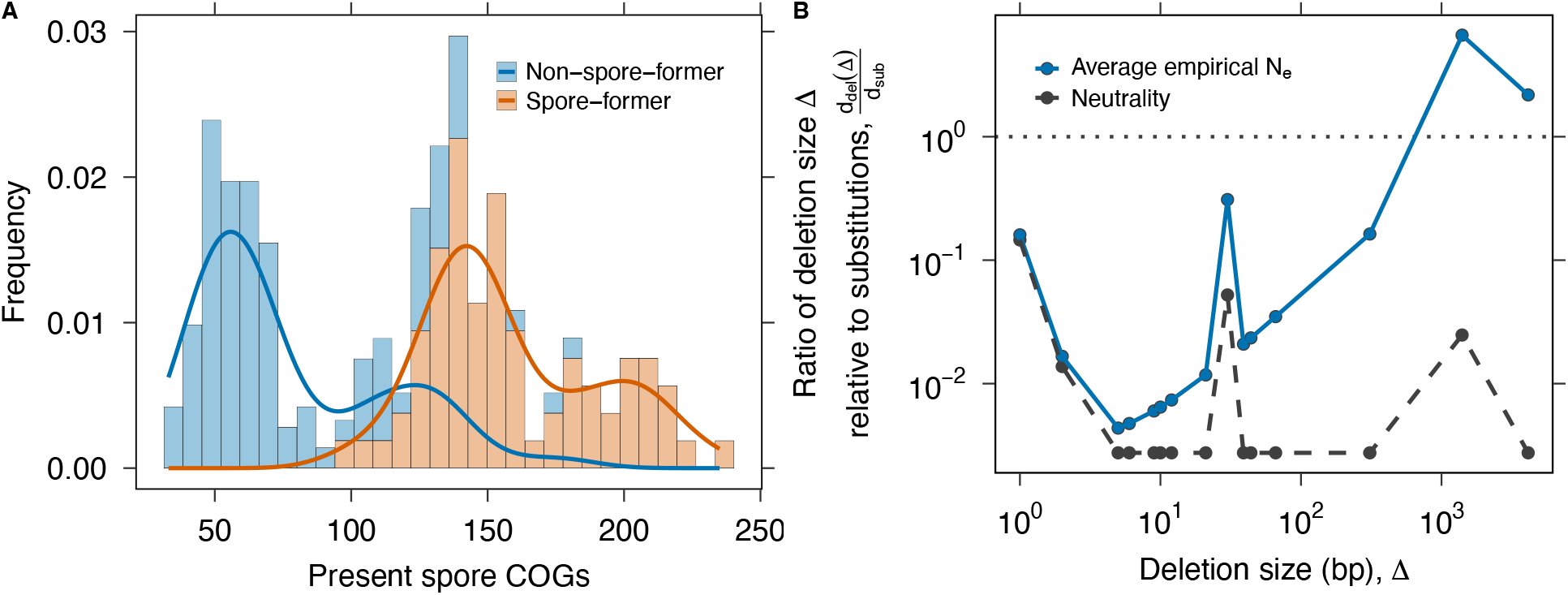
Evolutionary maintenance of sporulation. (A) Conservation of sporulation-related genes across spore-forming and non-spore-forming lineages in the Bacillota phylum based on the frequency of COGs (Clusters of Orthologous Genes). There are 237 COGs in total [15]. (B) The loss of sporulation genes in *Bacillus* can be examined by combining energetic cost estimates, which inform the strength of selection (*s*), with empirical mutation rates [76]. Here, we calculate the fixation rate of beneficial deletions relative to that of neutral substitutions (analogous to *dN/dS* in the population genetics literature; Supporting Information). The black dashed line shows the neutral expectation (*s* = 0), where the ratio is determined solely by the mutation rates of deletions and substitutions. The solid blue line shows the average non-neutral case using published estimates of *N*_*e*_ [5, 76]. For deletion sizes where the ratio is greater than 1, the energetic cost of maintaining sporulation genes is sufficient for selection to favor their loss. The sharp peaks reflect the empirical distribution of deletion sizes.

The loss of complex traits is a recurring theme in evolution, often attributed to relaxed selection and the accumulation of neutral mutations [2, 33]. Yet even in classic examples such as eye reduction in cave-dwelling fish or flight loss in birds, growing evidence suggests that the regression of biological function can be favored by selection, especially when a trait imposes physiological burdens or energetic costs [31, 86]. Our study provides a first-principles demonstration of how the energetic cost of a complex trait can influence its evolutionary fate. More broadly, the trade-offs we identify may apply to other metabolically expensive traits across the tree of life, including bioluminescence, motility, and secondary metabolite production in microbes, symbiotic nodulation and inducible defenses in plants, and elaborate courtship displays or environmentally contingent morphologies in animals. Although our model focuses on sporulation, a widespread form of dormancy that supports the persistence and dispersal of a globally abundant group of bacteria, the broader approach of quantifying energetic investment and linking it to evolutionary dynamics provides a scalable framework for understanding the distribution and loss of complex traits in nature.

## Methods

### Bioenergetic accounting: definitions and assumptions

We estimated bioenergetic costs using glucose as the sole carbon and energy source. In bacteria, glucose is metabolized through the tricarboxylic acid (TCA) cycle, yielding ATP. The hydrolysis of ATP to ADP and inorganic phosphate *(P*_*i*_) releases approximately 30 kJ/mol of free energy (Δ*G*) under standard conditions. While some ATP is used to power energy-consuming processes, other hydrolysis events support the biosynthesis of macromolecular building blocks. Following established conventions [43, 80], we estimate cellular energy expenditure in ATP (or ATP-equivalents) and use *P* to denote one hydrolysis event (GTP, etc., counted 1:1 as P). Following [43, 46], we partition costs into *direct* (*P*_*D*_) and *opportunity* (*P*_*O*_). Direct costs capture ATP-powered steps such as monomer activation/charging and polymerization (e.g., aminoacyl-tRNA charging, chain elongation). Opportunity costs capture the energetic value of precursor synthesis (e.g., NADPH-consuming reductions) that forgoes growth if diverted to a trait. Total cost is *P*_*T*_ = *P*_*D*_ + *P*_*O*_. We report *P*_*T*_ unless stated otherwise, because it best reflects resource allocation under balanced growth. However, *P*_*D*_ is useful when considering heat dissipation and instantaneous power, and *P*_*O*_ is useful when considering that some monomers can be recycled during processes. We assume 20°C for budget references and use the growth (*C*_*G*_) and maintenance (*C*_*M*_) entries of [43] where indicated.

It is important to note that biosynthetic costs vary across species due to differences in metabolic pathways and environmental factors such as resource availability [51]. Our analysis focused on cellular processes with sufficient quantitative data, including those related to the central dogma and membrane synthesis. We excluded minor costs below approximately 10^5^ ATP, such as post-translational modifications, and protein folding [43], although even these small expenditures may be evolutionarily significant in certain regulatory contexts [27].

#### Replication costs

Costs associated with the replication of the mother cell genome are incurred early in the spore cycle, before the commitment stage *∼* 2*h*. Although DNA replication is a complex process involving unwinding, primer synthesis, Okazaki fragment ligation, and proofreading, most of the energy is spent on synthesizing nucleotide building blocks and polymerizing them into DNA.

Let *L*_*g*_ be the genome length (bp). With 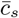 the *direct* synthesis cost of a DNA nucleotide (excluding polymerization; *≈* 11 *P*_*D*_), *c*_*p*_ the polymerization cost (2 *P*_*D*_*/*nucleotide), *c*_hel_ = 1 *P*_*D*_*/*bp for helicase unwinding, and *c*_prim_ = 0.32 *P*_*D*_ */*bp for lagging-strand primers [43, 46], we write:

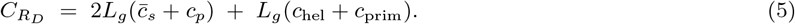

The *opportunity* cost uses the average opportunity per DNA nucleotide, 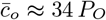 (precursor synthesis *∼* 33 *P*_*O*_ plus conversion), giving:

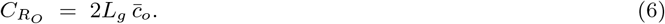

Ligation and proofreading are *<* 10^*−*4^ of 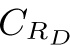 and are neglected.

#### Transcription costs

Transcription is also affected by complex mechanisms such as activation, initiation, termination, proofreading, and RNA processing. From a bioenergetic perspective, however, most of the transcription budget is dedicated to nucleotide synthesis and polymerization [43]. Therefore, the expression costs of genes can be estimated as the sum of the costs associated with protein-coding genes involved in spore formation and spore revival. Like replication, transcription costs have two components: direct costs associated with the polymerization and the opportunity costs, which are the energy needed to synthesize the ribonucleotide building blocks. Assuming that the length of genes and mRNAs (premature and mature) are approximately the same as in other bacteria [43], we estimate the direct costs of transcription as:

i. *One-time costs*: We assume efficient nucleotide recycling, so nucleotide synthesis for each transcript is charged once. Let *L*_mRNA,*j*_ be the length of transcript *j, N*_mRNA,*j*_ the number of transcripts produced over the program, and 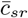 the direct synthesis cost per RNA nucleotide (i.e., RNA direct cost minus polymerization; 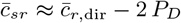). The corresponding opportunity cost uses 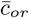 per nucleotide:

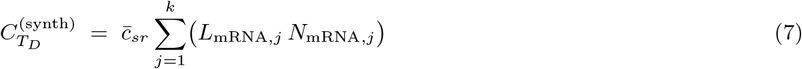

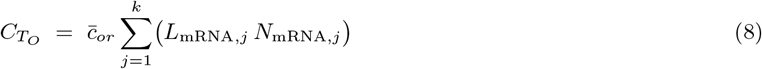 Here 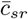 is the average direct costs of synthesizing an RNA nucleotide (10 *P*_*D*_), and 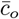 is the average opportunity cost of synthesizing an RNA nucleotide, which is *≈* 31.5 *P*_*O*_.
ii. *Time-resolved polymerization*: polymerization is paid whenever a transcript is made. We distribute the transcript production of gene *j* across hours *t* using weights *w*_*j,t*_ proportional to the measured expression at hour *t*, with Σ_*t*_ *w*_*j,t*_ = 1. If *R*_*j*_ (*t*) = *N*_mRNA,*j*_ *w*_*j,t*_ transcripts are made at hour *t*, the hourly direct cost is:

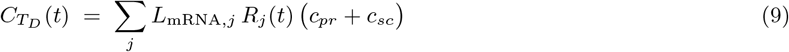

where *c*_*pr*_ = 2 *P*_*D*_ per nucleotide is the RNA polymerase elongation cost, and *c*_*sc*_ is a per nucleotide small direct overhead for relieving transcription-induced supercoils, motivated by the twin-domain model of Liu & Wang (1987) and the mechanochemistry of DNA gyrase (*≈* 2*ATP* per negative supercoil); we parameterize this as *c*_sc_ = 0.1 *P*_*D*_ */*nucleotide, typically *<*5% of polymerization [38]. The totals over the program are 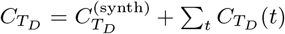 and 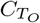 from Eq. 8.

To connect to protein demand, we estimate *N*_mRNA,*j*_ from protein abundances (parts per million, ppm) and a yield *Y* of proteins per mRNA lifetime: *P*_*j*_ = (ppm_*j*_/10^6^) *P*_total_, *N*_mRNA,*j*_ = *P*_*j*_*/Y*. We use *P*_total_ *≈* 1,774,445 proteins per cell [45] and a representative *Y* = 100. Because this yield-based accounting already fixes the number and timing of transcript synthesis via *w*_*j,t*_, we do not introduce an explicit mRNA degradation rate in the cost formulas to avoid double-counting.

#### Translational costs

Although mRNA has a shorter half-life and ribonucleotide polymerization is relatively expensive, proteins are 100-1000 times more abundant than transcripts [43]. In addition, nucleotides are already activated molecules, whereas tRNA must be charged with amino acids, making chain elongation more energetically demanding [43]. As a result, most of the energetic cost of protein synthesis is attributed to translation, while processes such as initiation, termination, and post-translational modification contribute minimally to total expenditure [43]. We estimated the per-cell translation costs as:

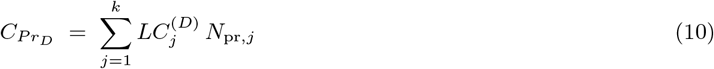

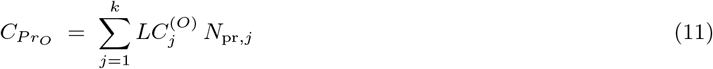

where 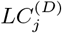 includes *∼* 4 *P*_*D*_ per incorporated amino acid for aminoacylation and elongation, and 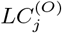 uses amino-acid opportunity costs (bacterial mean *∼* 25 *P*_*O*_) [43]. Protein degradation during the sporulation window is slow relative to synthesis, so we neglect turnover terms.

Beyond elongation and aminoacylation, we add a constant 2 *P*_*D*_ per completed protein to account for one GTP at initiation (IF2) and one GTP during termination/recycling (RF3), consistent with standard bacterial translation cycles [63]. Other small terms (e.g., proofreading, chaperone cycles) were not included, consistent with our *<* 10^5^ ATP cutoff.

#### Membrane synthesis and remodeling

We estimated the energetic cost of synthesizing and remodeling lipid bilayers for the cell envelope and sporulation-specific structures. The number of lipids is the bilayer area divided by the head-group area *a*_1_ = 0.65 nm^2^ [32, 54, 60]. We modeled the vegetative cell as a *spherocylinder* (cylinder + hemispherical caps) with length *L* = 2.5 *µ*m and diameter *D* = 1 *µ*m [3]. Writing *a* = *L/*2 and *b* = *D/*2, and using bilayer thickness *h* = 4 nm (3.0 nm hydrophobic core + 0.5 nm headgroup radius per leaflet) [37, 52], the outer and inner leaflet areas are:

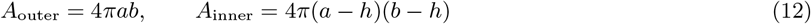

Assuming a membrane protein occupancy of *ϕ*_prot_ = 0.5, the lipid count is:

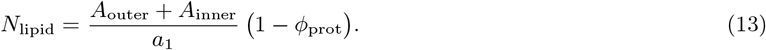

We take unit lipid costs as 18 *P*_*D*_ and 212 *P*_*O*_ [46], giving:

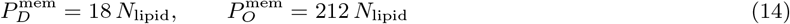

The septum is modeled as a flat circular bilayer (thickness effects negligible relative to *b*), with total bilayer area:

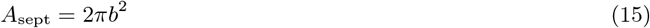

The costs follow analogously with the same *ϕ*_prot_ and unit costs.

During revival (0–3.5 h), we assume that only a fraction *f*_rev_ of the full vegetative membrane is newly synthesized and that a fraction *f*_recycle_ = 1*/*6 of lipid cost is offset by recycling. Net revival membrane cost is:

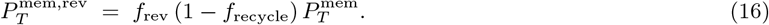

We use *f*_rev_ = 0.30 in the main analysis.

### Empirical data used for bioenergetic accounting

We used various publicly available cellular and molecular information to support this theoretical framework.

#### Gene sets and lengths

Gene and protein metadata (including lengths and SubtiWiki symbols) were obtained from SubtiWiki [90]. To unify identifiers, we normalized gene tokens and propagated locus tags across synonym lists.

#### Reference genome and counting conventions

All genome–level quantities were referenced to *Bacillus subtilis* subsp. *subtilis* strain 168 (NCBI RefSeq NC 000964.3; assembly GCF 000009045.1, ASM904v1). The genome length used in our calculations is *L*_*g*_ = 4,215,606 bp. Unless stated otherwise, gene counts refer to protein-coding loci (CDS); the total number used here is *n*_CDS_ = 4,237. Non-coding RNAs were excluded from gene-count totals but may appear in expression sets where relevant. All percentages “of the genome” and “of genes” are computed relative to these values.

#### Protein sequences and per-protein costs

Reviewed UniProt sequences for *B. subtilis* were mapped deterministically to locus tags via BSU identifiers in name/gene fields, then through unambiguous SubtiWiki primary/alias tokens. Per-protein direct and opportunity costs 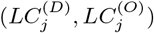 were computed by summing amino-acid costs from [46] over each sequence.

#### Protein abundances

PaxDb abundances (ppm) [81, 82] were tokenized and mapped to locus tags using the same symbol map. When a PaxDb row matched multiple tags, ppm was split evenly (mass-conserving). Proteins without a match received a tiny floor (0.1 ppm), then all ppm were renormalized to the known total protein count *P*_total_ = 1,774,445 [45] to obtain copies per cell.

#### Sporulation time series

Expression heatmaps from SporeWeb [13, 56] were converted to long format (*t*_0_–*t*_8_). For each gene *j*, hourly weights *w*_*j,t*_ *∝* normalized expression were used to (i) assign one-time transcript synthesis at first appearance and (ii) distribute polymerization across hours (Methods: *Transcription costs*).

#### Revival (germination/outgrowth)

Newly synthesized proteins from Swarge *et al*. [77] were converted to interval scores as successive differences (H0.25, H0.5, H1, H1.5, H2.5, H3.5). We excluded H5.5 (210–330 min) to avoid vegetative growth. For each protein, interval weights (normalized positive scores) distribute copies and thereby transcription/translation costs over time. Revival membrane remodeling used *f*_rev_ = 0.30 with *f*_recycle_ = 1*/*6 as above.

#### Handling missing fields

When gene or protein length was unavailable in SubtiWiki or the mapped UniProt entry, we substituted the dataset median (gene length and protein length medians computed over the available entries). If a protein lacked sequence-derived per-protein cost totals, we estimated them as (length × median per–amino-acid cost) computed across proteins; if length was also missing, we used protein-level medians. For proteins without a PaxDb match we assigned a small abundance floor (0.1 ppm) and then renormalized all ppm values to the total protein count, ensuring mass conservation.

#### Head-to-head bioenergetics

Gene sets for complex traits (biofilm, motility/chemotaxis, heat shock, competence, homeostasis) were taken from SubtiWiki and mapped to locus tags, then joined to absolute protein copies per cell and per-protein amino-acid costs. For each trait, *translation* cost was computed as copies × (per-protein direct + opportunity costs). *Transcription* cost was treated as a one-time mRNA synthesis sufficient to produce those proteins (transcripts = proteins/yield), split into direct (nucleotide synthesis; polymerization 2 ATP/nucleotide is *not* included for these build-only bars) and opportunity costs. Trait bars exclude maintenance/turnover as well as per-gene DNA replication and membrane synthesis; those appear once as global references to avoid double-counting.

The “spore life cycle” bar aggregates *build-only* transcription and translation for spore formation, germination (0–0.25 h), and outgrowth (0.5–3.5 h). “Maintenance during cycle” equals *C*_*M*_ (ATP·cell^*−*1^·h^*−*1^) multiplied by the program window (8 h + 0.25 h + 3.25 h). The “total cell budget” is *C*_*G*_ + *t*_gen_*C*_*M*_ for one generation (here *t*_gen_ = 1.16*h* at 20°C; *C*_*M*_ and *C*_*G*_ are 1.16 *×* 10^9^ and 9.25 *×* 10^10^ from [43]). The upper x-axis reports positions as a percentage of this budget. All quantities are per cell at 20*°*C; gene lists were curated to avoid overlap.

### A bioenergetic model of spore efficiency

To investigate how energy limitation influences sporulation efficiency, we developed a population dynamics model of *Bacillus* that incorporates the metabolic demands of endospore formation and vegetative cell production. We used bioenergetic yield parameters to capture how efficiently resources are converted into either spores or cells. The yields of vegetative cells (*Y*_*v*_) and spores (*Y*_*s*_) are defined as:

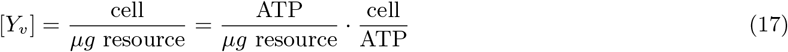

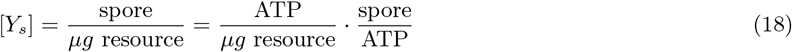

Using our ATP estimates, we can parameterize these yields:

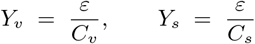

where *ϵ* represents the ATP produced per *µg* of resource, reflecting the efficiency of energy extraction from a given substrate. These expressions link the yields of cells and spores, reducing the degrees of freedom in our model by one. Given our definition of yield and the focus of our study on bioenergetics, it is important to explicitly account for the consumption of a limiting resource (*R*). To do so, we apply a minimal model of microbial growth in batch culture, defined as follows:

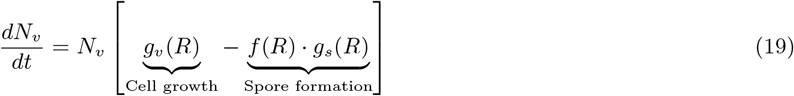

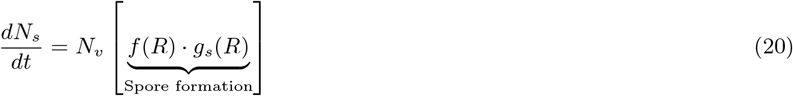

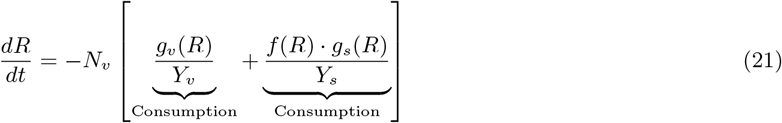

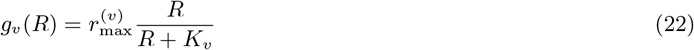

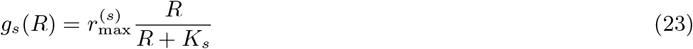

where *r*_*max*_ is the maximum growth rate, and *K* is the half-saturation constant, specifying the value of *R* where *g*(*R*) is half of *r*_*max*_. The rate of spore initiation is modeled as the following function:

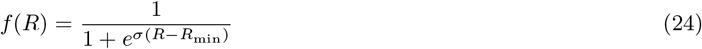

The sporulation initiation function exhibits intuitive properties. The sensitivity parameter *σ* governs the steepness of the transition from a low to high rate of spore formation. When *σ* ≫ 1, the function approximates a Heaviside step function [1], and can be defined as:

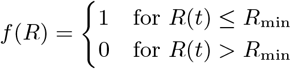

The inclusion of a spore germination term is unnecessary for a deterministic system where the resource concentration only decreases over time.

### Evolutionary maintenance of sporulation

In this section, we use a minimal model of natural selection acting on a single locus to illustrate the selective advantage conferred by the loss of a single nonfunctional nucleotide via a mutation-induced deletion. We focus on a scenario in which a population of spore-forming *Bacillus* inhabits an environment that supports continuous growth, making sporulation unnecessary. In this context, sporulation genes are not expressed and become effectively nonfunctional. As a result, they fall under relaxed selection [11], allowing mutations to accumulate over time, including those that would be deleterious in environments where sporulation is required.

However, nonfunctional DNA can still impose a metabolic cost. A cell that loses a single nucleotide through a mutation-induced deletion may gain a fitness advantage relative to its ancestor due to having been relieved of this slight, though existent, metabolic burden. This advantage can be calculated as the number of ATP required to build nucleotides (*c*_*n*_ *≈* 50 ATP), expressed relative to the cell’s total energetic budget [43]. In the case of *Bacillus*, the selective advantage of the mutant can be calculated as:

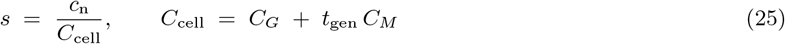

Under the strong-selection, weak-mutation evolutionary limit, the probability that a mutation of frequency *f* will fix can be derived as [10]:

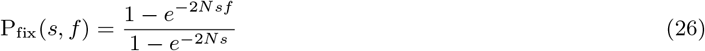

A *de novo* mutation has an initial frequency of 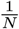, from which we obtain for *s* ≪ 1:

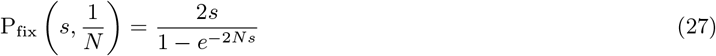

where *N* is the effective population size. Deletions are then successfully fixed at the following rate:

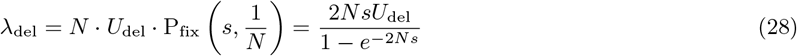

In the limit of an extended timescale with a low deletion rate, the rate of successful deletions can be approximated as the number of successful deletions [19].

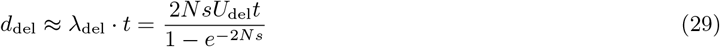

We have identified how positive selection can reduce the energetic demands imposed by a nonfunctional region of the genome. The function *d*_del_, however, is somewhat unwieldy because it depends on the number of elapsed generations. To simplify interpretation, it is useful to normalize this function by the rate of putatively neutral substitutions. Since deletions have been defined in terms of fitness gains, it is reasonable to treat substitutions at the same sites as neutral. Therefore, we consider only substitutions within the region encoding endospore formation as neutral. The rate of substitution in this region can be defined as:

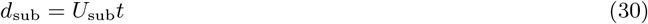

By taking the ratio, we can acquire a time-independent quantity:

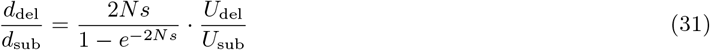

This equation is effectively the classic ratio of nonsynonymous to synonymous substitutions, adjusted for differing mutation rates. This adjustment is important because *U*_del_ is typically lower than *U*_sub_ [39, 40, 42, 76]. In *Bacillus*, mutation accumulation experiments estimated the rate of insertions and deletions at 1.20 *×* 10^*−*10^ per event, per individual, per generation, while the rate of substitution was 3.35 *×* 10^*−*10^, nearly three times higher [76]. By counting the fraction of indels that were identified as deletions in each reported mutation accumulation line, we can estimate a deletion rate of 6.0 *×* 10^*−*11^.

Using these rates, we evaluated the likelihood that a beneficial deletion in spore formation and revival genes becomes fixed relative to a neutral substitution, given the energetic cost of a single nucleotide. Under strict neutrality, this ratio is expected to be less than one due to the lower mutation rate of deletions (Fig. S3). To examine how this ratio scales with deletion size, we applied two published estimates of the effective population size for *Bacillus*: 6.119 *×* 10^7^ from [76] and 3.224 *×* 10^8^ from [5].

To better capture how metabolic costs influence the evolution of nonfunctional DNA in *Bacillus*, additional features can be incorporated into the model. For example, deletions do not always remove a single nucleotide. Large segments of the bacterial genome can be lost through individual deletion events [57]. We can define the strength of selection on a deletion of size Δ as the product of the deletion size and the relative metabolic burden of a single nucleotide, *s*_Δ_ = *s ·* Δ. However, these larger fitness gains may be rare due to the lower probability of observing large deletions [87].

We can examine the empirical probability distribution of Δ in *Bacillus*. Although it does not capture all possible deletion sizes, it reflects a general pattern supported by data (Fig. S3). Smaller deletions are more frequent than larger ones. We incorporated this distribution into our calculation of the ratio *d*_del_*/d*_sub_, defining a size-specific deletion function:

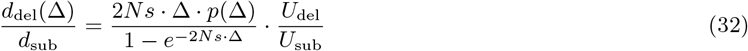

where *p*(Δ) is the empirical probability of acquiring a deletion of size Δ.

## Supporting information

suppplement

## Acknowledgments

We acknowledge support from the US National Science Foundation (DEB-1934554 and DBI-2022049 to JTL), the US Army Research Office Grant (W911NF2210014 to JTL), the National Aeronautics and Space Administration (80NSSC20K0618 to JTL), and the Alexander von Humboldt Foundation (to JTL). We thank DA Schwarz, BK Lehmkuhl, and J Ahmed for valuable discussions and technical assistance.

