## Supplementary material for "Evolutionary bioenergetics of sporulation": suppplement

### 1. Predicting sporulation efficiency in batch culture

In a batch culture design, we can operate under mass balance using the initial conditions of the system. Here, the sum of the supplied concentration of resources and the concentration of spore and vegetative cell abundances, corrected for yield, sets the total concentration of mass in the system. This sum must be equal to the sum at time $t$.

$$R_0 + \frac{N_v(0)}{Y_v} \equiv R(t) + \frac{N_v(t)}{Y_v} + \frac{N_s(t)}{Y_s} \tag{S1}$$

We have specified that no endospores were present at inoculation ($N_s(0) = 0$), and that all resources were supplied at the start of the experiment. These resources decrease in concentration as the concentration of cells and spores saturates, establishing a timescale $t_{sat}$ where the resources are effectively depleted $R(t_{sat}) \approx 0$. At this timescale, the mass balance relation reduces to the following:

$$R_0 + \frac{N_v(0)}{Y_v} \equiv \frac{N_v(t_{\mathrm{sat}})}{Y_v} + \frac{N_s(t_{\mathrm{sat}})}{Y_s} \tag{S2}$$

from which we can rearrange to obtain functions of the abundances of vegetative cells and spores.

$$N_v^* \equiv N_v(t_{sat}) = Y_v \left[ R_0 + \frac{N_v(0)}{Y_v} - \frac{N_s^*}{Y_s} \right] \tag{S3}$$

$$N_s^* \equiv N_s(t_{sat}) = Y_s \left[ R_0 + \frac{N_v(0)}{Y_v} - \frac{N_v^*}{Y_v} \right] \tag{S4}$$

Using these stationary values, we can derive a prediction for the fraction of spores in the population, known as sporulation efficiency:

$$\begin{aligned}\phi &\equiv \frac{N_s^*}{N_s^* + N_v^*} \\ &= \left[1 + \frac{N_v^*}{N_s^*}\right]^{-1} \\ &= \left[1 + \frac{Y_v}{N_s^*}\left(R_0 + \frac{N_v(0)}{Y_v} - \frac{N_s^*}{Y_s}\right)\right]^{-1}\end{aligned} \tag{S5}$$

So, we have a prediction of sporulation efficiency that only depends on one observable, the stationary density of spores $N_s^*$. An equivalent prediction can be obtained that only relies on $N_s^*$.

We can obtain a form of this result that only depends of parameters by considering 1) the growth regime $R_0 \ll K_v$, $K_s$, linearizing the growth functions $g(R)_v$, $g(R)_s$ and 2) the $\sigma \gg 1$ limit of $f(R)$. In this parameter regime spores do not form over timescale $t_{\mathrm{min}}$, reducing the system to

$$\frac{dN_v}{dt} = N_v R \frac{r_{\mathrm{max}}^v}{K_v} \tag{S6}$$

$$\frac{dR}{dt} = -N_v R \frac{r_{\mathrm{max}}^v}{K_v Y_v} \tag{S7}$$

Over timescale $t_{\mathrm{min}}$ the mass balance equation is then

$$R_0 + \frac{N_v(0)}{Y_v} \equiv R(t) + \frac{N_v(t)}{Y_v} \tag{S8}$$

Solving for $N_v(t)$, we remove the dependency on $R(t)$ and obtain

$$\begin{aligned}\frac{dN_v}{dt} &= N_v\left(R_0 + \frac{N_v(0)}{Y_v} - \frac{N_v(t)}{Y_v}\right)\frac{r_{\mathrm{max}}^v}{K_v} \\ &= N_v \tilde{r}\left(1 - \frac{N_v}{\tilde{K}}\right)\end{aligned} \tag{S9}$$

where we have defined $\tilde{r} \equiv \frac{r_{\mathrm{max}}^v}{K_v}\left(R_0 + \frac{N_v(0)}{Y_v}\right)$ and $\tilde{K} \equiv Y\left(R_0 + \frac{N_v(0)}{Y_v}\right)$ as the effective growth rate and carrying capacity. We recognize this as the ordinary differential equation (ODE) for logistic growth, the solution of which is:

$$N_v(t) = \frac{\tilde{K}}{1 + \left(\frac{\tilde{K} - N_v(0)}{N_v(0)}\right)e^{-\tilde{r}t}} \tag{S10}$$

from which we obtain the number of vegetative cells when sporulation initiates, $N_v(t_{\mathrm{min}})$. After $t_{\mathrm{min}}$, the dynamics of the system are modeled as follows:

$$\frac{dN_v}{dt} = N_v R\left(\frac{r_{\mathrm{max}}^v}{K_v} - \frac{r_{\mathrm{max}}^s}{K_s}\right) \tag{S11}$$

$$\frac{dN_s}{dt} = N_v R \frac{r_{\mathrm{max}}^s}{K_s} \tag{S12}$$

$$\frac{dR}{dt} = -N_v R\left(\frac{r_{\mathrm{max}}^v}{K_v Y_v} + \frac{r_{\mathrm{max}}^s}{K_s Y_s}\right) \tag{S13}$$

If we assume that the growth of vegetative cells is negligible after $t_{\min}$, then $N(t_{\min}) \approx N_v(t_{\text{sat}})$, removing a state variable from our spore efficiency solution. This result provides an example of how sporulation efficiency could be predicted under batch culture conditions.

### 2. Predicting sporulation efficiency in a chemostat

As a point of comparison, it is worth contrasting the above result with the solution one obtains by using a chemostat instead of a batch culture. This can be accomplished by amending each equation in our batch culture system with 1) a dilution term $\delta$ that defines the fraction of total vessel volume that flows in and out of the chemostat per unit time and 2) the concentration of resources supplied to the vessel $R_0$.

$$\frac{dN_v}{dt} = N_v \left[ \underbrace{g_v(R)}_{\text{Cell growth}} - \underbrace{f(R) \cdot g_s(R)}_{\text{Spore formation}} - \underbrace{\delta}_{\text{Dilution}} \right] \tag{S14}$$

$$\frac{dN_s}{dt} = N_v \underbrace{f(R) \cdot g_s(R)}_{\text{Spore formation}} - N_s \underbrace{\delta}_{\text{Dilution}} \tag{S15}$$

$$\frac{dR}{dt} = \underbrace{\delta R_0}_{\text{Input}} - \underbrace{\frac{N_v g_v(R)}{Y_v}}_{\text{Consumption}} - \underbrace{\frac{N_v f(R) \cdot g_s(R)}{Y_s}}_{\text{Consumption}} - \underbrace{\delta R}_{\text{Dilution}} \tag{S16}$$

This model marks a clear and biologically meaningful difference from a model of batch culture dynamics, as there now exists a stationary non-zero resource concentration ($R^* > 0$). The absence of a term describing the process of germination is justified if the rate of germination per-unit spore concentration is much larger than $\delta$. The stationary solution can be obtained by setting the derivatives to zero.

$$\begin{aligned} 0 &= g_v(R^*) - f(R^*) g_s(R^*) - \delta \\ 0 &= N_v^* f(R^*) g_s(R^*) - N_s^* \delta \\ 0 &= \delta R_0 - \delta R^* - \frac{N_v^* g_v(R^*)}{Y_v} - \frac{N_v^* f(R^*) \cdot g_s(R^*)}{Y_s} \end{aligned} \tag{S17}$$

We can make analytic progress by assuming that the stationary resource concentration is less than the half-saturation constants, $R^* \ll K_v, K_s$, a limit that can be experimentally obtained by tuning the dilution rate. This limit allows us to make the linear approximation $g(R^*) \approx \frac{R^* r_{\max}}{K}$. Furthermore, if $R^* \ll R_{\min}$, we can use the Heaviside limit for the rate of spore formation. Using these approximations, we obtain the following stationary solutions:

$$N_v^* = \left[ \frac{r_{\max}^{(v)}}{K_v Y_v} - \frac{r_{\max}^{(s)}}{K_s Y_s} \right]^{-1} \cdot \left[ R_0 \left( \frac{r_{\max}^{(v)}}{K_v} - \frac{r_{\max}^{(s)}}{K_s} \right) - \delta \right] \tag{S18}$$

$$N_s^* = \frac{N_v^*}{\delta} \left[ \frac{r_{\max}^{(v)} K_s}{r_{\max}^{(s)} K_v} - 1 \right]^{-1} \tag{S19}$$

$$R^* = \delta \left[ \frac{r_{\max}^{(v)}}{K_v} - \frac{r_{\max}^{(s)}}{K_s} \right]^{-1} \tag{S20}$$

From which we obtain sporulation efficiency:

$$\phi \equiv \frac{N_s^*}{N_s^* + N_v^*} \approx \frac{r_{\max}^{(s)} K_v}{r_{\max}^{(v)} K_s} \tag{S21}$$

where we have neglected dilution. This result can be interpreted as the ratio of the rates of cellular growth and endospore formation, with yield playing no role. This result means that one would expect energetic costs not to factor into sporulation efficiency, contrasting with the batch culture scenario. This analytic result is consistent with prior experimental efforts to quantify efficiency in a chemostat setting (Fig.S1; [1]).

### 3. Simulating batch culture dynamics

We investigated the validity of our sporulation efficiency predictions across spore formation parameter regimes, focusing on the two terms governing spore formation: the Monod function governing the formation of spores $g_s(R)$ and the rate of initiating spore formation $f(R)$. Our system of equations was solved using the `solve_ivp()` function from SciPy v1.10.1 for a given parameter combination [2], from which sporulation efficiency was calculated.

We found that our predictions tended to fail when the Monod constant is small relative to the concentration of supplied resources (Fig.S2), reducing the Monod function to the constant $r_{\max}^{(s)}$. This deviation is consistent with the assumptions in our derivation, as the limit $K_s \ll R_0$ effectively decouples resource consumption from growth. In contrast, our predictions remained consistently accurate across spore formation initiation parameters (Fig.S2). This result suggests that details of spore formation initiation may not considerably shape predictions obtained at an extended timescale.

**Table S1.** Parameter values used in simulations

| **Variable** | **Meaning** | **Value** |
|---|---|---|
| $R_0$ | Initial resource concentration | $10^3$ $\mu g$/mL |
| $N_v(0)$ | Initial cell density | $10^5$ cells/mL |
| $N_s(0)$ | Initial spore density | 0 spores/mL |
| $r_{\max}(v)$ | Maximum growth rate of cells | 2 $\text{hr}^{-1}$ |
| $r_{\max}(s)$ | Maximum rate of spore formation | 1/8 $\text{hr}^{-1}$ |
| $K_v$ | Monod constant for cellular growth | $3.2 \cdot R_0$ $\mu g$/mL |
| $K_s$ | Monod constant for spore formation | $0.1 \cdot R_0$ $\mu g$/mL |
| $\sigma$ | Sharpness of spore formation initiation | 0.1 mL/$\mu g$ |
| $R_{\min}$ | Resource threshold for sporulation | 5 $\mu g$/mL |
| $Y_v$ | Cell yield | $\epsilon(5 \cdot 10^{-11})$ cells/$\mu g$ |
| $Y_s$ | Spore yield | $\epsilon(1 \cdot 10^{-11})$ spores/$\mu g$ |

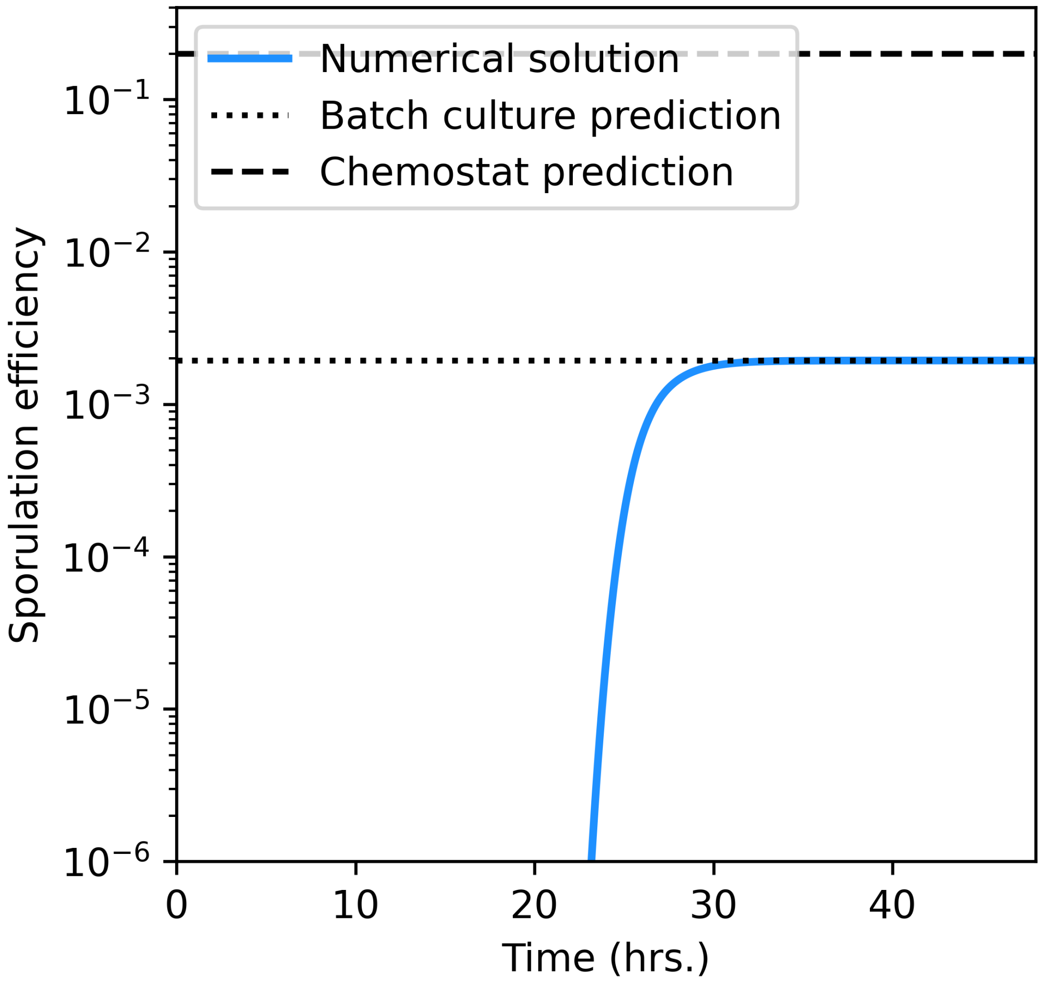


**Figure S1.** Our batch culture predictions of sporulation efficiency were validated by obtaining the numerical solution of our system of differential equations. As resources become depleted, predictions that depend on the yields of cells and spores (dashed line) become increasingly accurate. The prediction in the chemostat limit is plotted as a point of comparison.

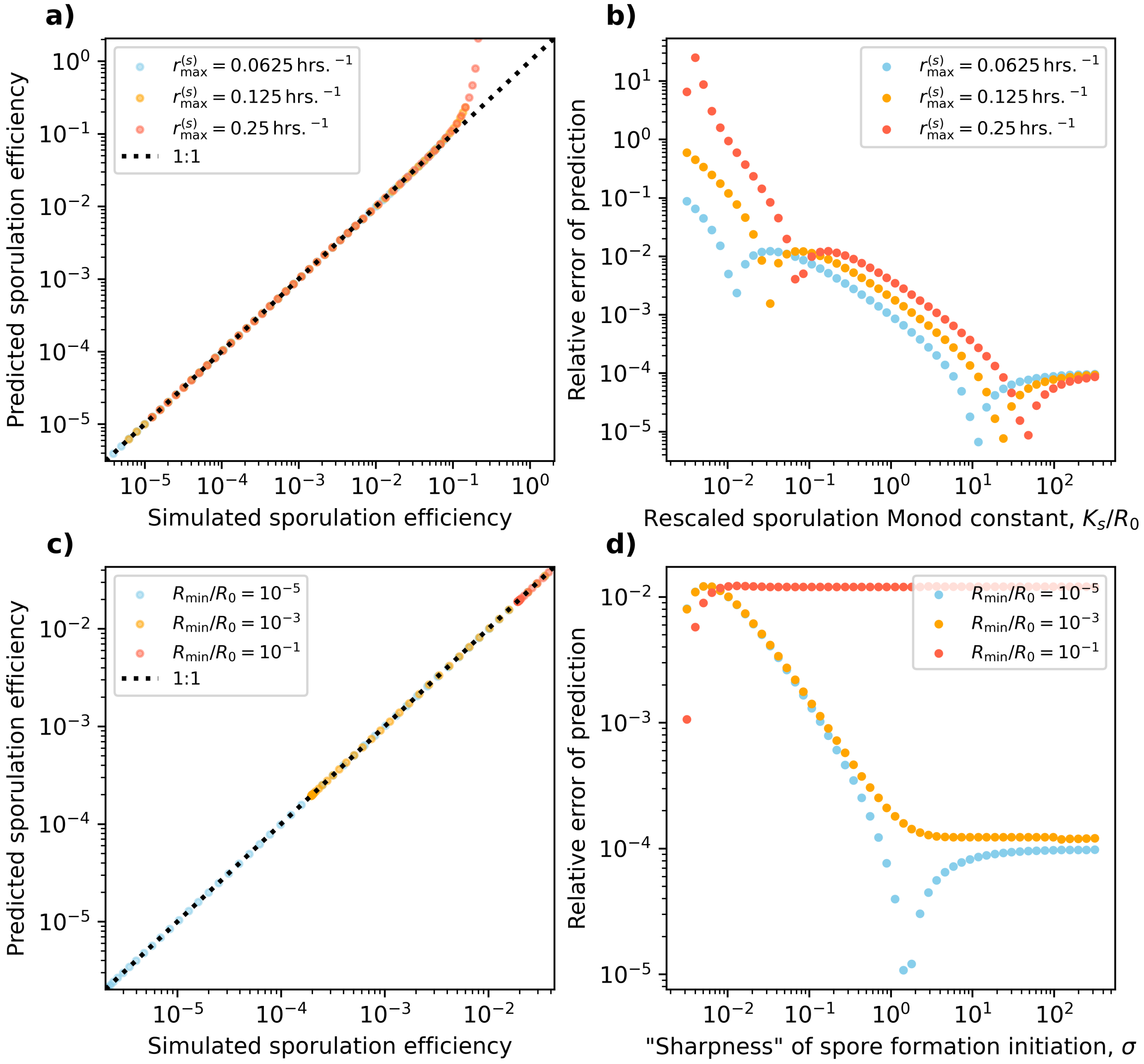


**Figure S2.** The effect of parameters controlling spore formation on the validity of sporulation efficiency predictions.

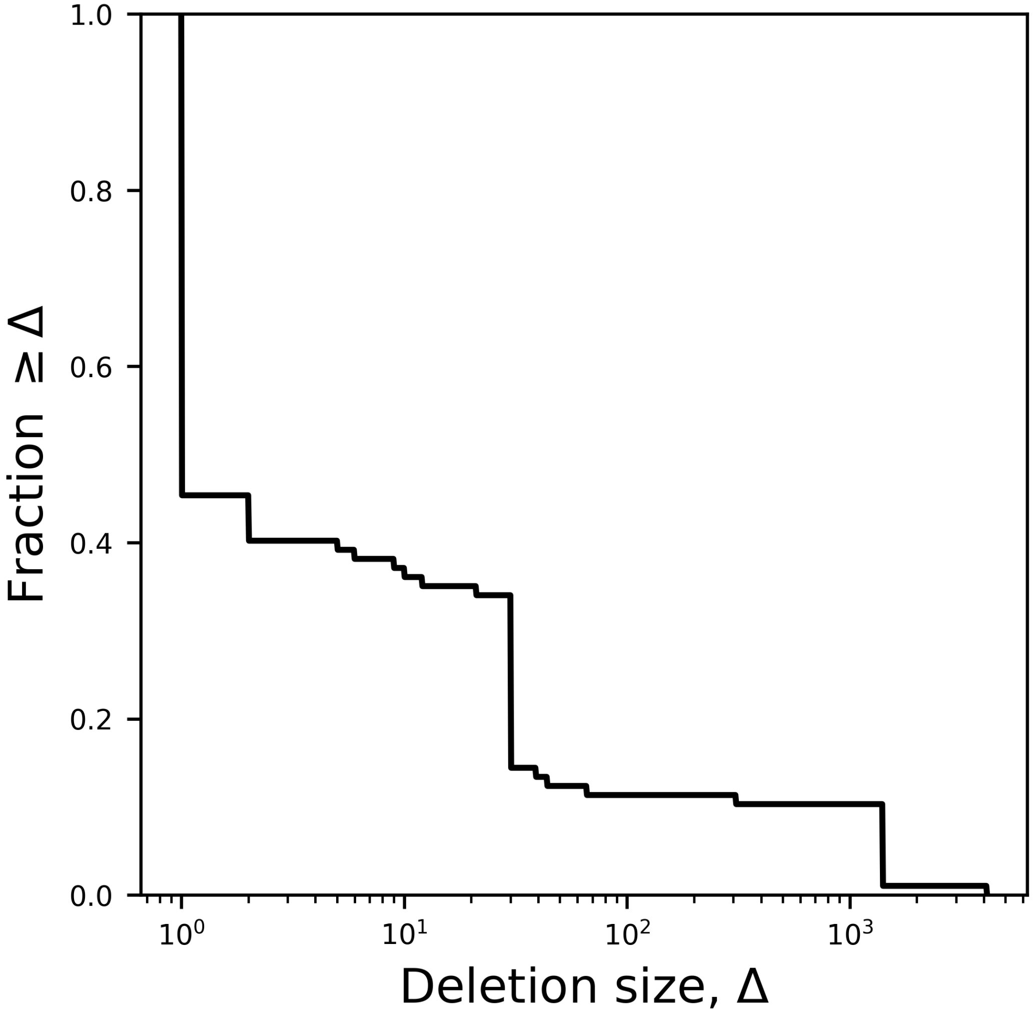


**Figure S3.** The empirical distribution of deletion sizes in *Bacillus subtilis*.

**Table S2.** Reported sporulation efficiencies from the literature.

| Publication | Efficiencies (%) | DOI |
|---|---|---|
| Grossman and Losick 1988 | 23, 34 | doi.org/10.1073/pnas.85.12.4369 |
| Kudo and Horikoshi 1979 | 0, 1, 5, 10, 15, 30, 35, 50, 55, 70, 95 | doi.org/10.1080/00021369.1979.10863868 |
| Monterio et al. 2008 | 10, 15.3, 16.7, 17, 23.6, 25.6, 40.9, 45.2, 45.9, 46.4, 47.9, 48, 48.8, 50.7, 53.2, 58.8 | doi.org/10.1021/bp050062z |
| Tavares et al. 2013 | 13.2, 15.1, 45.5, 74.2 | doi.org/10.1007/s00284-012-0269-2 |
| Widderich et al. 2015 | 0, 0.5, 1, 40, 70, 90 | doi.org/10.1111/mmi.13304 |
| Mendez et al. 2004 | 0.0025, 0.003, 0.0079, 0.014, 0.0213, 0.1 | doi.org/10.1128/JB.186.4.989-1000.2004 |
